# Rapid Therapeutic Recommendations in the Context of a Global Public Health Crisis using Translational Bioinformatics Approaches: A proof-of-concept study using Nipah Virus Infection

**DOI:** 10.1101/333021

**Authors:** Khader Shameer, Kipp W. Johnson, Ben Readhead, Benjamin S. Glicksberg, Claire McCallum, Amjesh Revikumar, Jamie S. Hirsch, Kevin Bock, John Chelico, Negin Hajizadeh, Michael Oppenheim, Joel T. Dudley

**Affiliations:** Centre for Research Informatics and Innovation; Northwell Health, New Hyde Park, NY, USA; Department of Information Services; Northwell Health, New Hyde Park, NY, USA; Institute for Next Generation Healthcare, Mount Sinai Health System, NY, USA; Institute for Computational Health Sciences, University of California, San Francisco CA, USA; Institutes of Health and Wellbeing, School of Computing Science at Glasgow, University of Glasgow, UK; Department of biotechnology and microbiology, Kannur University, Kerala, India; Department of Medicine, Division of Pulmonary Critical Care and Sleep Medicine, Center for Health Innovations and Outcomes Research (CHIOR), Northwell Health, NY; Division of Infectious Diseases, Zucker School of Medicine at Hofstra/Northwell, Hempstead NY

## Abstract

We live in a world of emerging new diseases and old diseases resurging in more aggressive forms. Drug development by pharmaceutical companies is a market-driven and costly endeavor, and thus it is often a challenge when drugs are needed for diseases endemic only to certain regions or which affect only a few patients. However, biomedical open data is accessible and reusable for reanalysis and generation of a new hypotheses and discovery. In this study, we leverage biomedical data and tools to analyze available data on Nipah Virus (NiV) infection. NiV infection is an emerging zoonosis that is transmissible to humans and is associated with high mortality rates. In this study, explored the application of computational drug repositioning and chemogenomic enrichment analyses using host transcriptome data to match drugs that could reverse the virus-induced gene signature. We performed analyses using two gene signatures: i) A previously published gene signature (*n*=34), and ii) a gene signature generated using the characteristic direction method (*n*= 5,533). Our predictive framework suggests that several drugs including FDA approved therapies like beclometasone, trihexyphenidyl, S-propranolol etc. could modulate the NiV infection induced gene signatures in endothelial cells. A target specific analysis of CXCL10 also suggests the potential application of Eldelumab, an investigative therapy for Crohn’s disease and ulcerative colitis, as a putative candidate for drug repositioning. To conclude, we also discuss challenges and opportunities in clinical trials (n-of-1 and adaptive trials) for repositioned drugs. Further follow-up studies including biochemical assays and clinical trials are required to identify effective therapies for clinical use. Our proof-of-concept study highlights that translational bioinformatics methods including gene expression analyses and computational drug repositioning could augment epidemiological investigations in the context of an emerging disease with no effective treatment.

## Introduction

Nipah Virus (NiV) is a member of *Paramyxoviridae* family of enveloped, Negative-sense single-stranded RNA viruses. NiV is a member of Henipavirus genus – other Henipaviruses include *Cedar henipavirus, Ghanaian bat henipavirus, Hendra henipavirus* and *Mojiang henipavirus* [1–9]. NiV infection is considered as an emerging infectious disease threat by the World Health Organization (WHO) (See: http://www.who.int/csr/disease/nipah/en/) [10, 11]. NiV infection was first reported and the virus was isolated during an outbreak in Malaysia and Singapore in the late 1990s [12–14]. NiV infection reemerged in 2018 in Kozhikode, a coastal city in Kerala, India. The Kerala reemergence is associated with high mortality rate (above 80% of identified cases, personal communication) (See: http://www.bbc.com/news/world-asia-india-44193145). At present there are no approved therapies or vaccines against NiV. In this study, we explore the application of translational bioinformatics databases, tools and methods like computational drug repositioning to predict potential FDA-approved, therapies using publicly available datasets. Follow-up studies, including experimental assays and clinical trials are required before using repositioned candidate drugs for clinical use.

### Clinical manifestations of NiV infection

NiV infection is characterized by a combination of neurological, respiratory and cardiovascular complications. These include but are not limited to: high fever, seizures, encephalitis, meningitis, tremor, ptosis, dysarthria, dysphasia, respiratory tract lesions, severe acute respiratory distress syndrome (ARDS), tachycardia, myocarditis, vomiting, hypertension, and segmental myoclonus along with or without other clinical features [15–23]. The incubation period for the NiV infection is 4 to 14 days with a combination of clinical symptoms, although some infected patients may remain asymptomatic [24, 25].

### Genome, proteome and host-virus interaction studies

The genome sequence of the NiV was reported with 18246 nucleotides that encode viral proteins (See: http://www.uniprot.org/uniprot/?query=proteomeUP000128950) including the nucleocapsid protein, phosphoprotein, matrix protein, fusion protein, glycoprotein and RNA polymerase [26]. Mechanistic studies suggest several viral proteins function to target host proteins, for example phosphoproteins P/V/W/C interact and inactivate the transcription factor *STAT1* by inhibiting interferon-induced tyrosine phosphorylation [27–35]. Functional genomics studies have shown that micoRNAs like mir-181 could also play a key role in the NiV infection. Mir-181 is a key modulator of human immune function and neuroinfllammation and may play role in neurovirulence of NiV including blood-brain barrier disruption [36–40]. NiV M proteins also target *ANP32B*, an anti-apoptotic, phosphoprotein [41]. *APAF1* an apoptosis regulator has also shown role in the pathogenicity of NiV [42]. Functional receptors of NiV include tyrosine kinases like Ephrin-B2 and ephrin-B3 [43, 44]. Several comparative genome analyses are also reported for the viral genome and zoonotic reservoirs like bats [45–52].

### Candidate therapies and vaccines targeting NiV

Several antibody, drug and vaccine discovery efforts to target NiV infection are in progress for NiV [53–57]. Multiple vaccine development projects against NiV based on viral epitopes and host-virus interaction mechanisms are currently available for livestock animals [32, 58–98]. Notably, vaccine candidates like human monoclonal antibody m102.4 has found to be effective in pre-clinical studies [71]. Pre-clinical immunotherapy studies demonstrate that monoclonal antibodies might be beneficial (anti-G and anti-F MAbs) as agents against NiV infection. Small molecules that activate *IRF3* and modulate *RIG-I*-like receptors pathways [56] were also investigated as potential strategies to target NiV infection [54]. Drugs like ribavirin (a broad spectrum antiviral effective against both RNA and DNA viruses) have been shown to be associated with lower mortality rates, but lack conclusive evidence from randomized controlled clinical trials [53, 99–103]. Evidence from a recent studies suggests that Favipiravir (T-705), an investigational treatment for influenza may prevent NiV infection in a hamster model [104]. Efforts are also underway to develop nucleic acids therapeutics against NiV infection [105]. Most of these therapies need full cycles of clinical trials or additional evidences including comparative effectiveness to understand optimal contributions to outcomes [53].

### Drug repositioning – systematic search for compounds to combat NiV infection

Drug repositioning (or drug repurposing; See Figure 1) is a drug development strategy designed to reduce the time to develop and market a drug for diseases with no approved drugs or diseases that may need better therapy [106, 107]. Rare, orphan, endemic or neglected diseases may not be ideal areas of research and development for pharmaceutical companies. However, drug repositioning offers an alternate path in such settings. For example, National Institute of Health’s – National Center for Advancing Translational Sciences recommends that 7,000 rare and neglected diseases that currently lack effective treatments could benefit from drug repositioning compared to traditional drug discovery that costs billions of dollars and may take more than a decade to deliver a drug to the market. Multiple reports on computational, pre-clinical, off-validation, and other mode-of-success for drug repositioning are available in biomedical literature. But there was no comprehensive, integrative analysis of reported repositioned drugs, their indications, repositioned drug targets, or chemical properties of repositioned drugs. Recently, we compiled drug repositioning investigations from literature and other public data repositories and developed, the first comprehensive resource of drug repurposing (See RepurposeDB: A reference database of drug repositioning investigations - http,//repurposedb.dudleylab.org/). By integrating large-scale data on drugs, indication, side effects, and mechanism of actions, the computational drug repositioning process can be automated. Predictive data can be used for prioritizing candidate drugs for downstream studies including biochemical validation and clinical trials. Thus, computational drug repositioning offers a predictive framework for accelerated therapeutic recommendation in scenarios that requires immediate therapeutic stratification [106–114].

**Figure 1:**
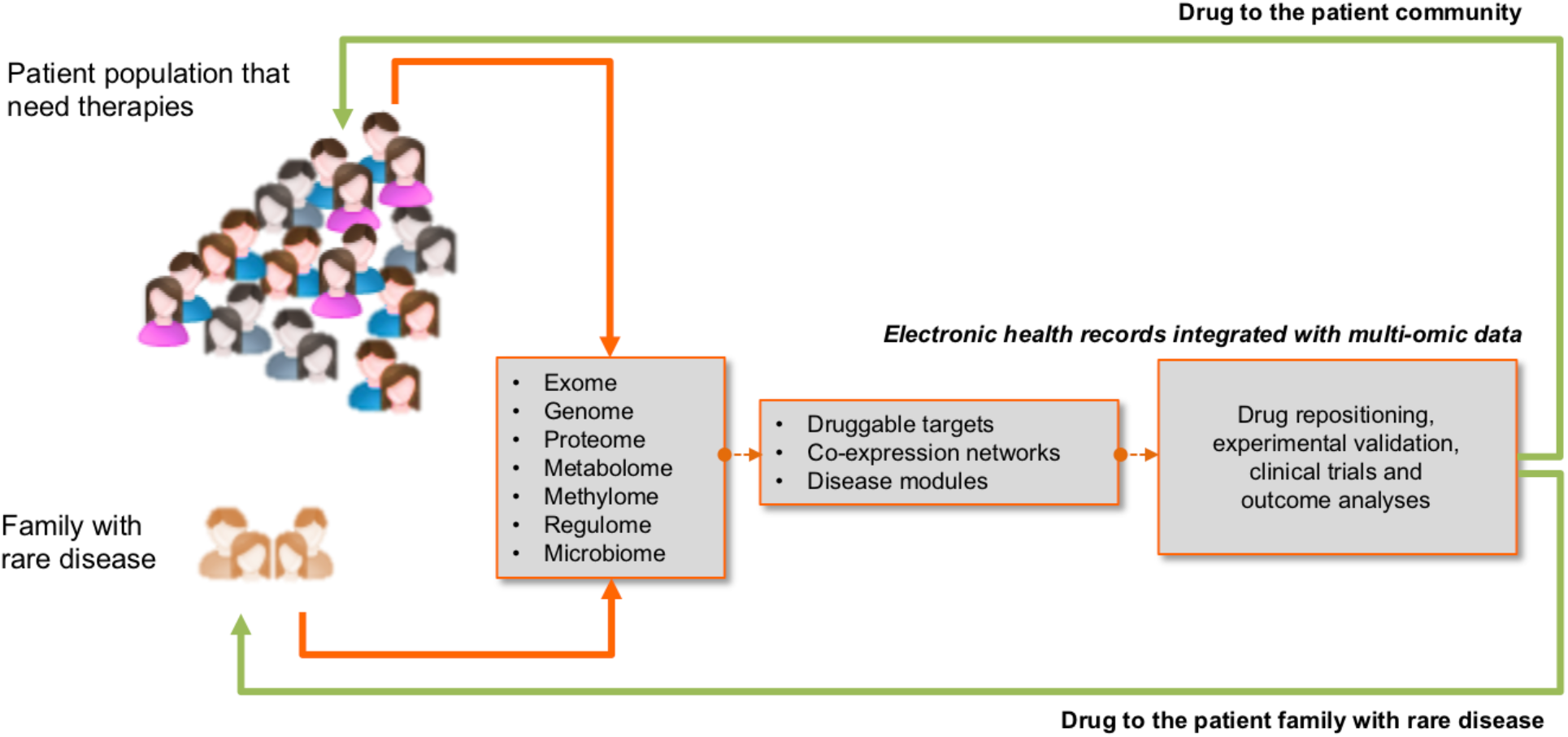
Concept of molecular-profiling based drug repositioning. Adaptation from an earlier version of a figure published in Shameer et.al, 2015 [107]

### Rationale for Computational Drug Repositioning to identify therapeutic indications for NiV infection

In the absence of an effective antiviral agent or prophylactic vaccine for NiV in human, it is imperative to develop better therapeutic agents to address such infectious disease threats. However, pharmaceutical companies have limited commercial prospects in developing drugs for rare, orphan, or neglected infectious diseases endemic diseases (Figure 2). Data-driven methods that combine computational and experimental approaches could complement, improve or reduce the cost of drug discovery. For example, during the first decade (1998-2008) of the NiV cases reported since its identification, the disease has an average case fatality rate of 52% over a total of 477 positively identified cases (using RT-PCR assays). (See: http://www.who.int/blueprint/priority-diseases/key-action/nipah/en/). In this context, leveraging drug repositioning may be beneficial. Drug repositioning could bring therapies to market in approximately half the budget and time required by traditional drug development cycle. Conducting drug repositioning analyses using publicly available translational bioinformatics tools, databases and methods may aid in such discovery to address public health crises and accelerate the path to discovery of new therapies. The American Medical Informatics Association defines “Translational Bioinformatics” as “the development of storage, analytic, and interpretive methods to optimize the transformation of increasingly voluminous biomedical data, and genomic data, into proactive, predictive, preventive, and participatory health”[115, 116]. In this study we show the application of a translational bioinformatics approach “computational drug repositioning” to find potential therapies to target NiV using publicly available datasets and tools. Therapeutic options to target this epidemiological threat is limited and drug repositioning will be a viable therapeutic identification strategy. However, since late ‘90s biomedical researchers have generated and deposited a variety of biomolecular data related to NiV in publicly accessible biomolecular databases.

**Figure 2:**
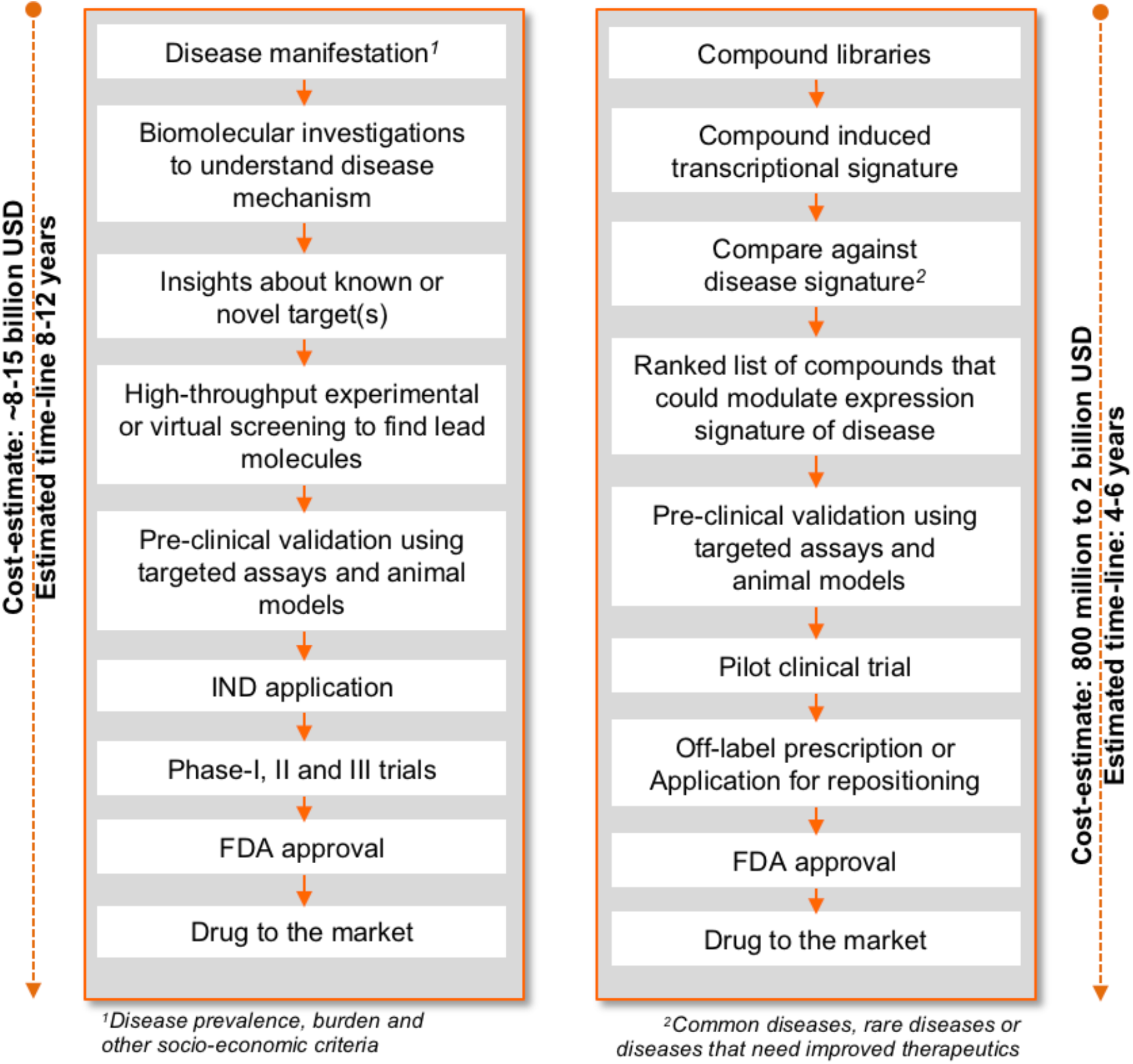
From medicine to markets: comparison of traditional drug discovery and drug repositioning. Steps in pipelines, cost-estimate and time-line from initial findings to the market from *Tobnick*, 2009 [184]. An earlier version of this figure was published in Shameer et.al, 2015 [107]

## Methods

Figure 3 illustrates the methodology workflow used in the present study. Methods include the following components: Aggregating NiV infection gene signature data for drug repositioning and Computational drug repositioning using NiV gene signatures followed by filtering and interpretation.

**Figure 3:**
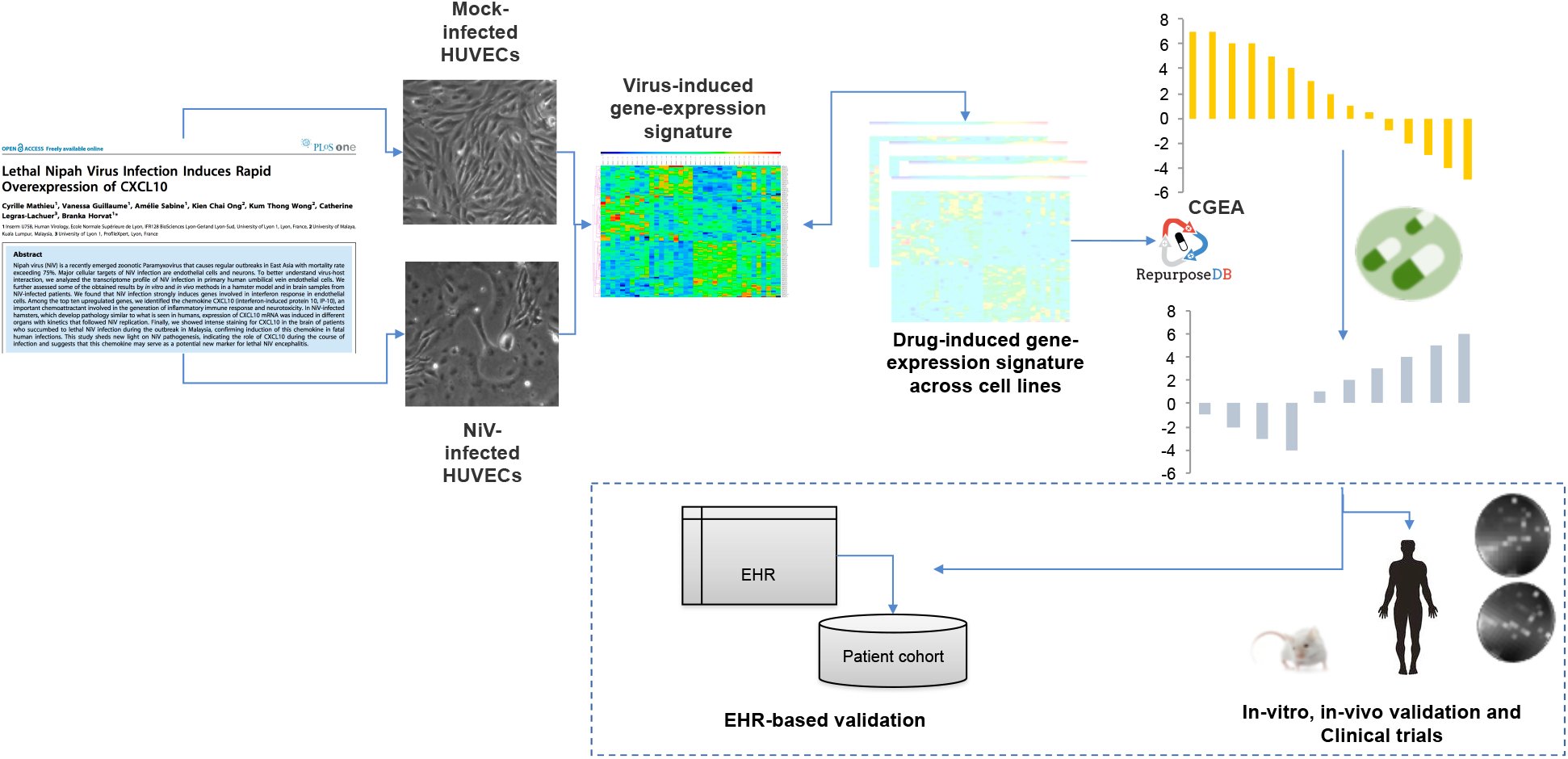
Computational drug repositioning workflow. Steps included in the dashed box illustrate potential future directions for the project.

### Aggregating NiV infection gene signature data for drug repositioning

A critical requirement of computational drug repositioning is the availability of gene expression datasets related to the disease of interest. We queried the Gene Expression Omnibus (GEO) [117] to collect NiV related gene expression data. We used published gene signature and used the published signature and a computed signature from the raw data (GEO accession code GSE33133). The samples were originally hybridized and analyzed using Codelink Uniset Human Whole Genome bioarrays and differential expression was originally computed using the Gene Spring v7.0 suite from Agilent [118]. After obtaining gene expression signatures differing between NiV-infected and control cells, we used Chemogenomic enrichment analyses (CGEA) method and evaluated drugs reported to reverse expression differences between NiV and control cells at an FDR-significance level of 0.10. We tested the gene signature against the signature of 1309 compounds, 743 of which have approval status in one or more global pharmaceutical markets as indicated in DrugBank See: https://www.drugbank.ca/drugs).

### Computational drug repositioning using NiV gene signatures

To leverage CGEA method, we first used the signature induced by NiV on endothelial cell lines, in the form of “upregulated” and “downregulated” gene identifiers. The input query of gene lists is matched to compounds, and the connectivity between the gene signature and compound is scored after various filtering steps against the available drug-induced signatures compiled from various reference databases. References resources like RepurposeDB (http,//repurposedb.dudleylab.org/), Connectivity Map (Cmap: https,//portals.broadinstitute.org/cmap/), Genomics of Drug Sensitivity in Cancer (GDSC: http://www.cancerrxgene.org/) or Cancer Cell Line Encyclopedia (CCLE: https,//portals.broadinstitute.org/ccle) are used to identify compounds that concordantly modulate the query signature in a direction “towards” or “away” from the input query. A Gaussian mixture model is used to derive the “connectivity score,” and statistical significance is estimated using the Kolmogorov-Smirnov test with the Benjamini-Hochberg method for control of false discovery rate. CGEA produces an output of a ranked list of candidate compounds that may potentially modulate a biological state of interest. Depending on the query signature and reference databases, many candidate compounds will often be extracted – such lists can be trimmed and prioritized for most likely candidates using annotations from reference databases (for example RepurposeDB, KEGG drugs, DrugBank, etc.) and also using specific characteristics including mechanism of action, side effects, chemical properties or biological targets. Detailed account of related methods to match gene signature to corresponding drugs are explained elsewhere [111, 112, 114].

## Results

### Compiling data sets for computational drug repositioning

Gene expression data was compiled from GEO (Samples retrieved on 20^th^ May 2018 https://www.ncbi.nlm.nih.gov/gds/?term=nipah%20virus). The data represents total of six samples (2 NiV infected (isolate UMMC1, Genebank AY029767) HUVEC cell lines with multiplicity of infection; 2 cell lines infected with recombinant viruses lacking the expression of accessory NiV C protein (NiVΔC) and 2 mock-infected HUVEC cells. Pathology studies indicate that the endothelium of central nervous system was susceptible to NiV infection and may thus represent a representative model system to study NiV infection using molecular level data [119].

i. *Published gene signature:* Published gene signature consists of 34 genes (upregulated genes: *n*=31; downregulated genes: *n*=3).
ii. *Characteristic direction-based gene signature:* Upregulated genes: *n*=2664; Downregulated genes: *n*=2869.

Eight upregulated genes were shared across both signatures (*IFIT1, IFI44L, OASL, CXCL10, IFIT2, OAS2, OAS1, IFI44*) and one gene was common across down regulated genes (*TAF4B*). Both gene lists were used independently to perform drug matching using CGEA approach using a library of drugs were annotated in conjunction with RepurposeDB. In the interpretation step, we primarily focus on the approved subset of compounds to extent the feasibility of launching immediate clinical trials. Briefly the gene set-drug matching data was compiled and annotated using data from the Connectivity Map, Anatomical Therapeutic Chemical (ATC) Classification System, PubChem, SIDER, Offsides and Drug Bank. Results compiled from CGEA consist of compound information including chemoinformatics signatures, drug target information, indications, mechanism of action and side effects.

### Computational Drug Repositioning using Gene Expression Signature Identifies Potential, FDA approved Therapies for Nipah Virus

Following sections summarize a subset of compounds predicted using computational drug repositioning. These drugs are not ready for clinical use and need extensive testing and clinical trials before use at the point of care.

### Published gene signature

Seven compounds have shown to optimally perturb the published gene signature (trihexyphenidyl, 5186223, convolamine, 5186324, tiletamine, S-propranolol). Two of these drugs are FDA approved for different indications, trihexyphenidyl (Score= −0.47, *P*=0.0006; parkinsonian disorders, drug-induced extrapyramidal movement disorders and antispasmodic drugs. See: https://www.drugbank.ca/drugs/DB00376); S-propranolol (Score= −0.44, *P*= 0.002) indications in acute myocardial infarction, arrhythmias, angina pectoris, hypertension, hypertensive emergencies, hyperthyroidism, migraine, pheochromocytoma, menopause, and anxiety; See: https://www.drugbank.ca/drugs/DB00571). Table 1 lists drugs capable of perturbing published gene signature with FDR <=0.10 (See: Supplementary Material for full list of compounds and annotations).

**Table 1:**
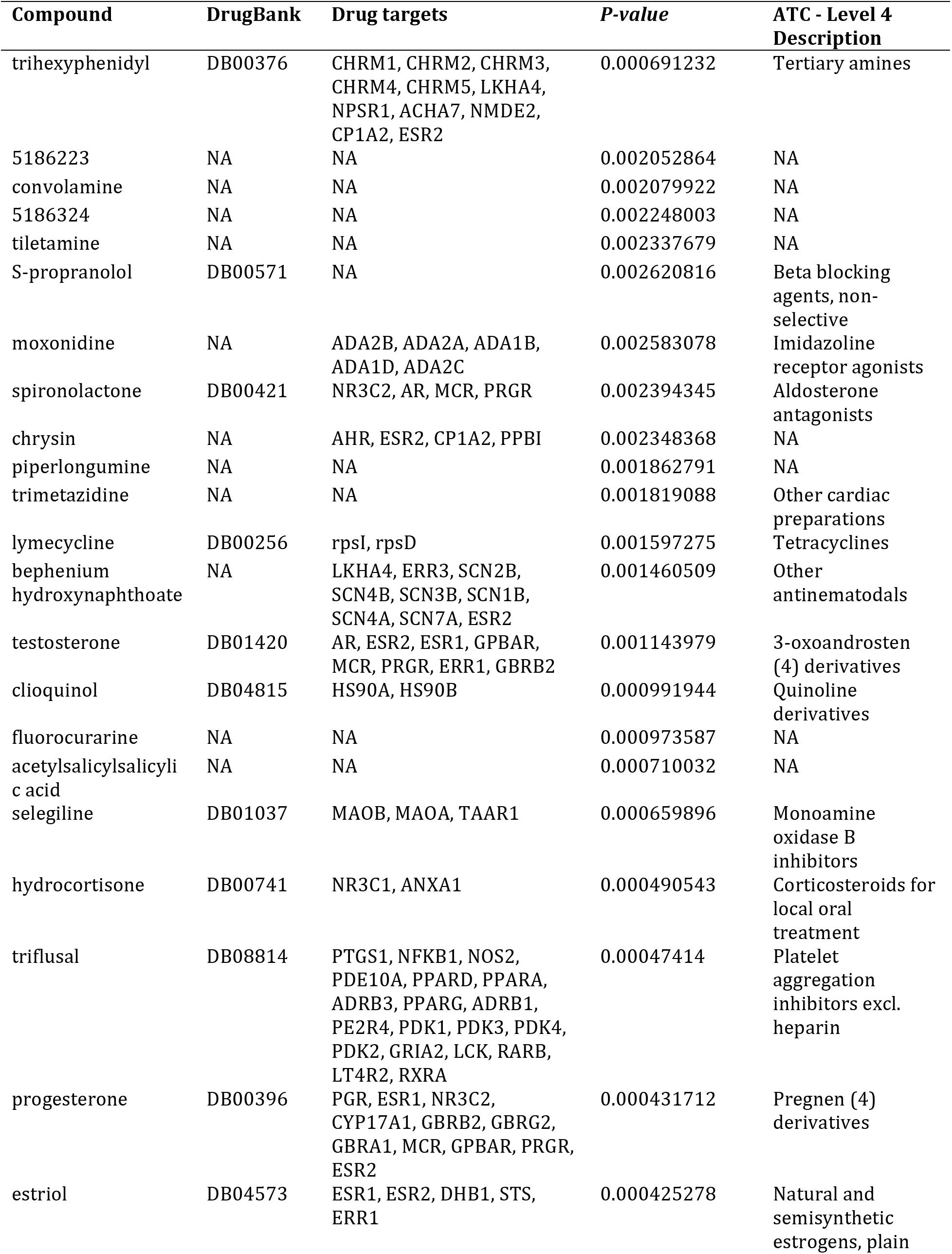
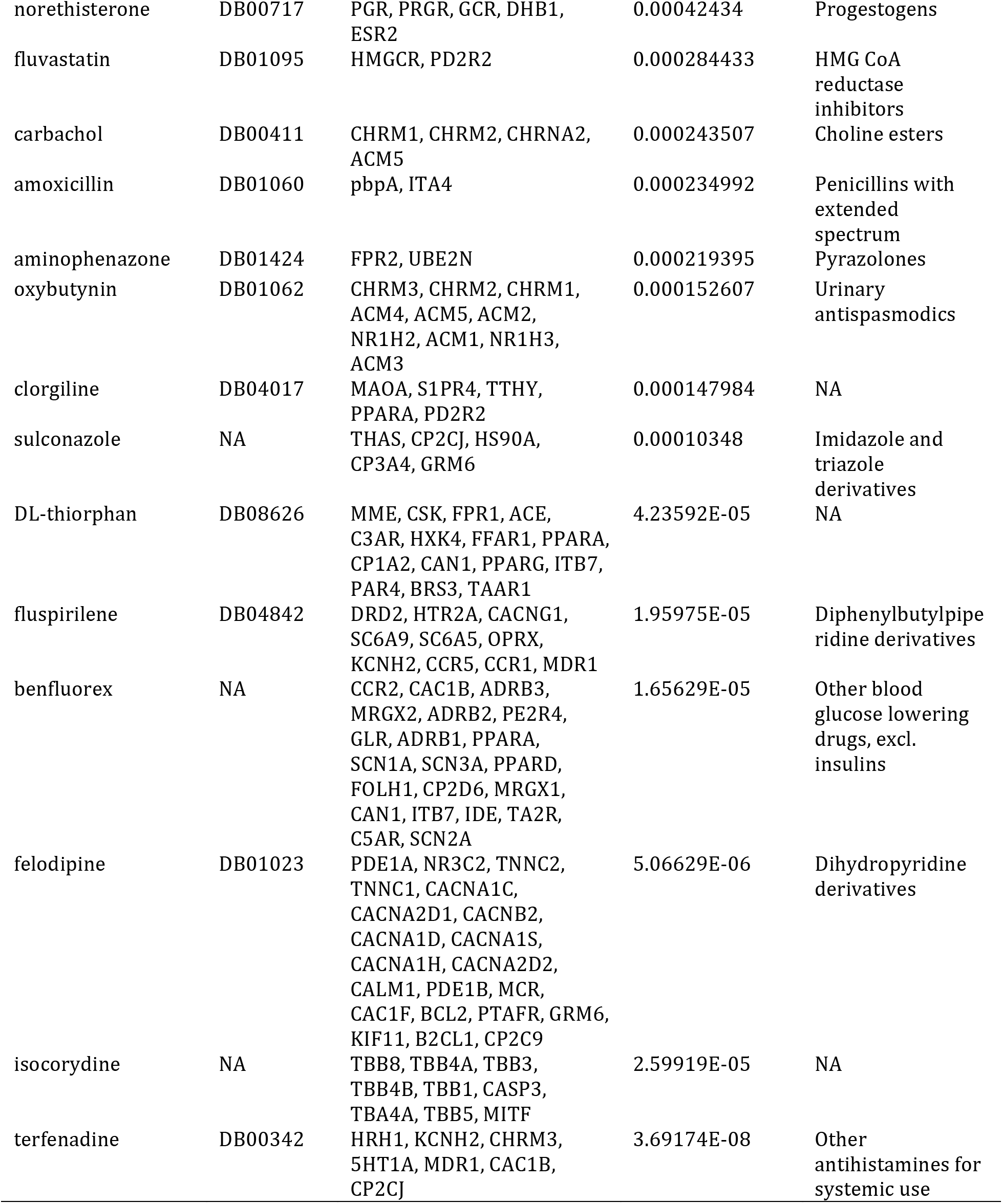
Drugs predicted to perturb the published gene signature

### Characteristic direction-based gene signature

Nine compounds have shown to optimally perturb Characteristic direction-based gene signature (5666823, 2-deoxy-D-glucose, oligomycin, pirinixic acid, clofilium tosylate, cantharidin, 0173570-0000 and beclometasone). Among these compounds, beclometasone (Score= −0.05, *P*= 0.001; See: https://www.drugbank.ca/drugs/DB00394) is an approved, anti-inflammatory small molecule with indications for asthma and allergic rhinitis (seasonal and perennial). It is currently an investigational compound for indications including Crohn’s disease and graft-versus-host disease. Table 2 lists drugs capable of perturbing characteristic direction gene signature with FDR <=0.10 (Also See Supplementary Material for full list of compounds and annotations).

**Table 2.**
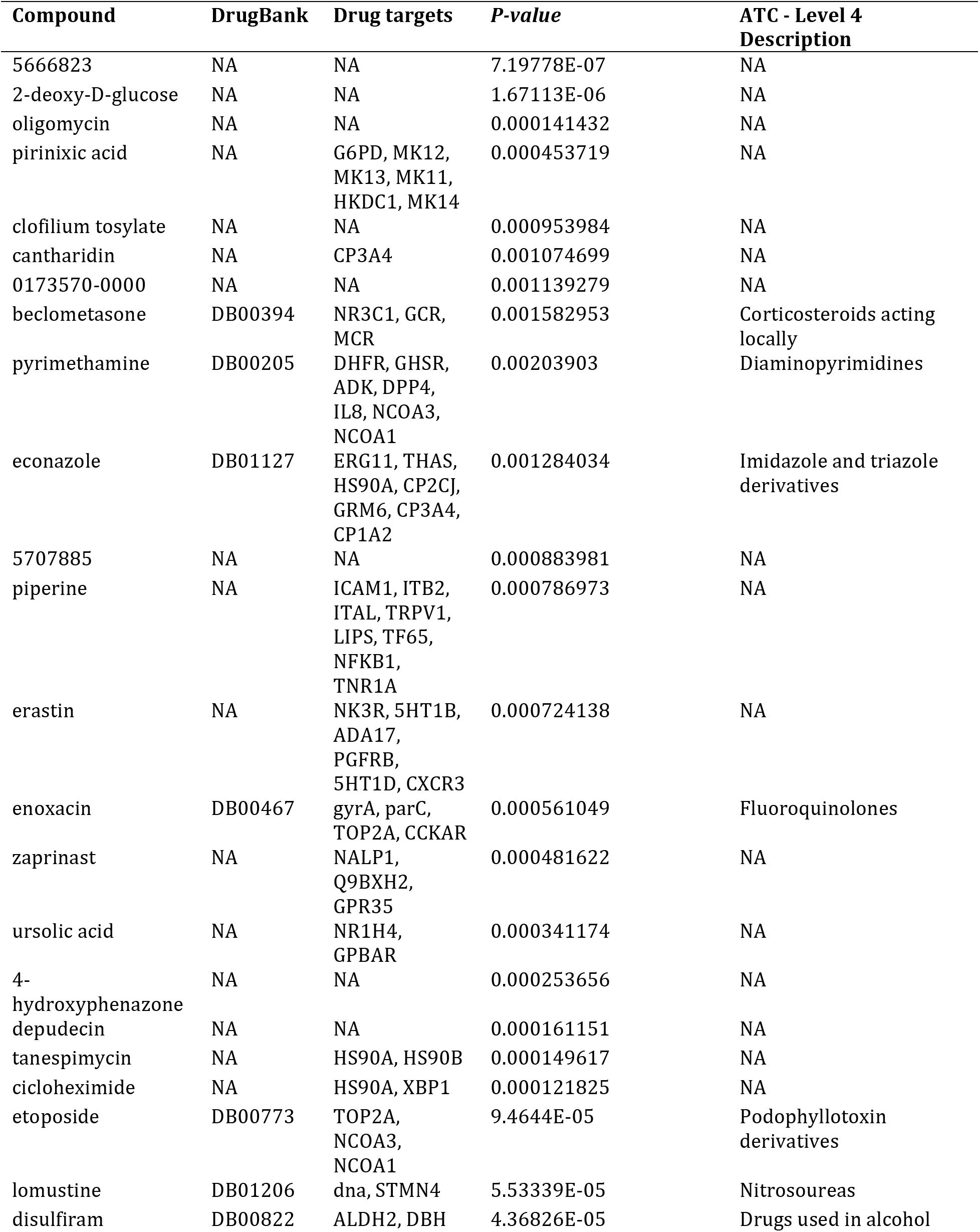
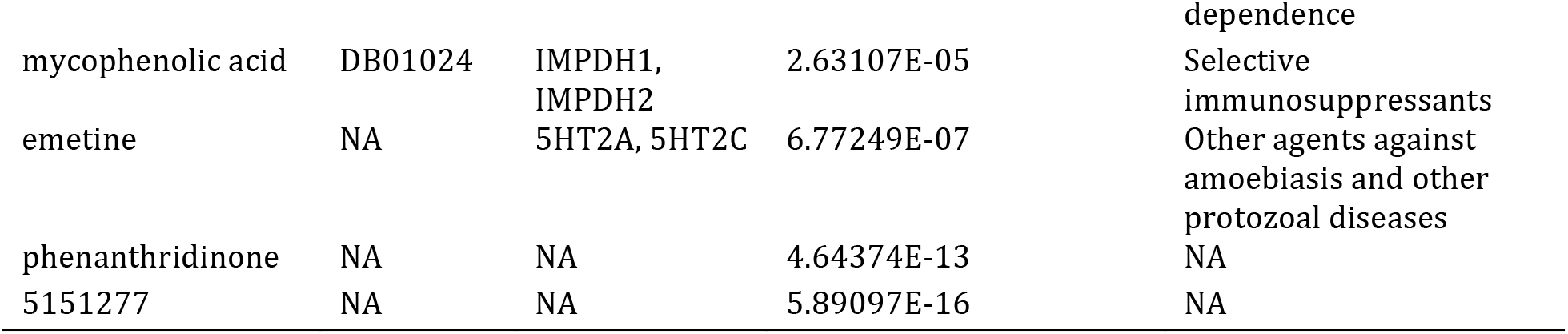
Drugs predicted to perturb the gene signature derived using characteristic direction method

### Antiviral candidate drugs from repositioning

Seven antiviral agents were available in our dataset. The following is the most effective to least effective order in published gene signature (ribavirin, vidarabine, moroxydine, zidovudine, saquinavir, zalcitabine, ganciclovir). While several of them had scores indicating activity but were not significant after FDR corrections. Ribavirin had the lowest score indicating most potent activity. Following was the order of activity in characteristic direction based gene signature: saquinavir, ganciclovir, moroxydine, vidarabine, zalcitabine, zidovudine, ribavirin. Ribavirin was the least effective - this is an example how the genes themselves, number of genes, and signature directionality of genes could influence the ranking of compounds. We also noted that moroxydine had optimal direction of score in both gene sets but were not statistically significant. Moroxydine is an antiviral agent structurally similar to ribavirin [120]. Conserved structural moieties often indicate similar chemical activities. Additional studies are required to elucidate the role of moroxydine as an anti-NiV agent.

### Targeting CXCL10 - a hallmark of NiV infection in humans

C-X-C motif chemokine (with synonyms C7; IFI10; INP10; IP-10; crg-2; mob-1; SCYB10; gIP-10) is an antimicrobial gene that encodes a chemokine of the CXC subfamily and ligand for the receptor CXCR3. CXCL10 was a gene expressed across both signatures (Figure 4(a)) and has a broad expression across 14 different tissue types including appendix and lymph node [121]. CXCL10 is involved in multiple inflammatory diseases (rheumatoid arthritis, inflammatory bowel disease and multiple sclerosis) that affect different systems of human physiology (See: https://www.ncbi.nlm.nih.gov/gene?db=gene&report=generif&term=3627) [121, 122]. CXCL10 also plays a key roles in several infectious diseases including chronic hepatitis B, tuberculosis etc. [122]. While no small molecules could target CXCL10, a fully human antibody (Eldelumab) that targets CXCL10 is reported. The molecule is currently under investigation for Ulcerative colitis [123, 124]. CXCL10 is a modulator of cytokine interaction networks and implicated in pathways including Chemokine Signalling pathway, TNF signalling pathway, Toll-like receptor signaling pathway, NOD-like receptor signalling pathway, NF-kappa B signalling pathway, RIG-I-like receptor signalling pathway. CXCL10 is also a member of disease pathways including rheumatoid arthritis, legionellosis, helicobacter pylori infection, salmonella infection, influenza A, and pertussis (Figure 4(b)).

**Figure 4:**
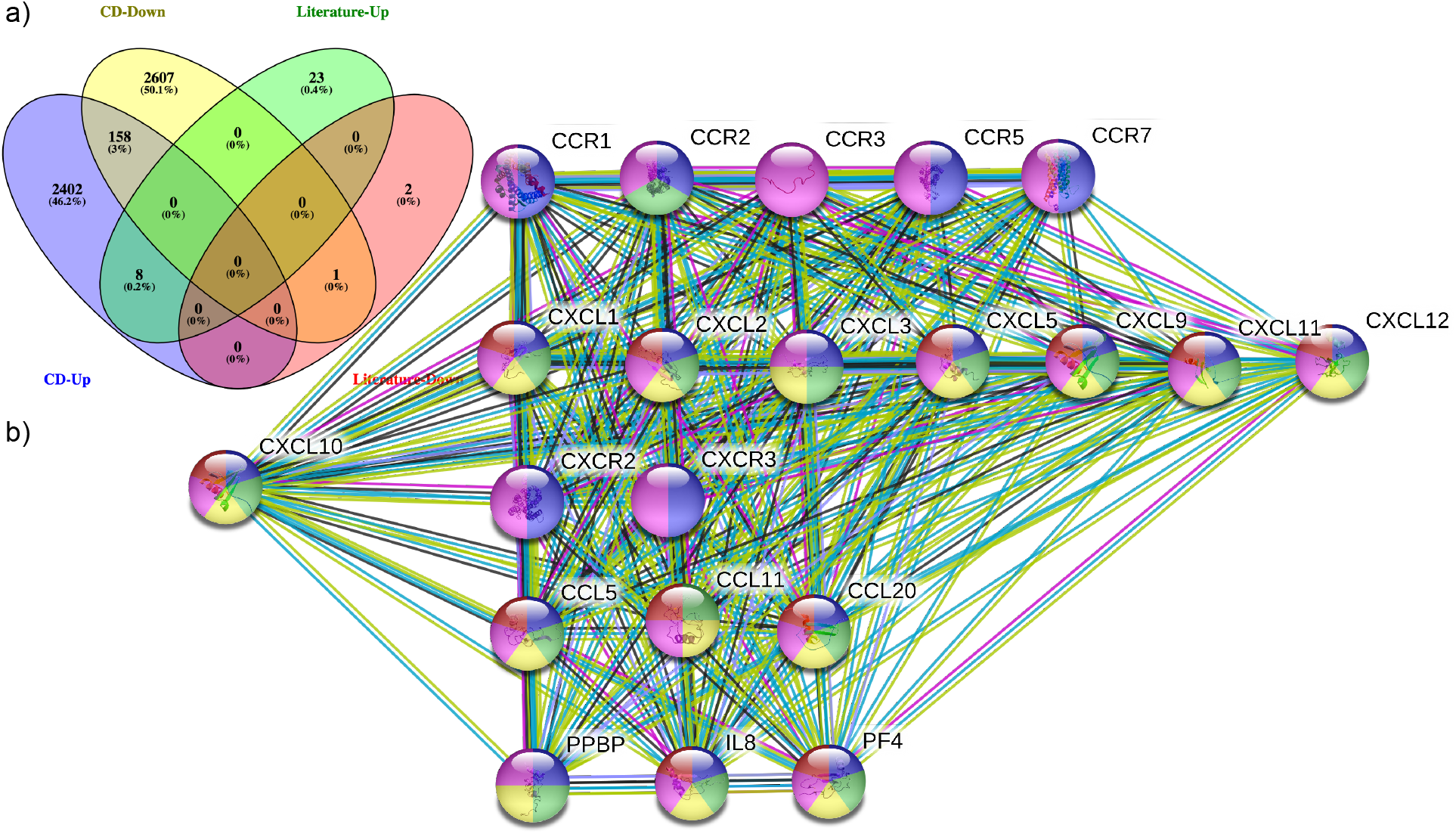
**a)** Overlap between signatures and direction of expression **b)** First degree interactome of CXCL10. Colors indicates various biochemical functional roles inferred from gene set enrichment analyses of interacting partners (PPI enrichment *P*=< 1.0e-16) of CXCL10 (blue: cell chemotaxis (biological process, *n*=19, FDR= 3.03e-35; green: chemokine receptor binding (molecular function, *n*=15, FDR=1.42e-32; **yellow**: extracellular space (cellular component, *n*=14; FDR= 6.4e-10); **magenta**: Cytokine-cytokine receptor interaction (KEGG pathways, *n*=21; FDR=9.72e-39); **red**: Small cytokines (intecrine/chemokine), interleukin-8 like (Pfam domains, *n*=12, FDR= 5.06e-29).

## Discussion

Precision medicine aims to leverage molecular profiling data to recommend medications based on an individual patient’s risk profile, existing medications, comorbidities, and non-clinical factors like diet and environmental exposure [125]. Precision medicine approaches that leverage mutation profiles to recommend therapies are emerging as a standard of care in oncology and potentially expanding to other therapeutic domains including cardiology [126]. Our current study provides, a hypothesis generating, proof-of-concept study to leverage such molecular profile drug stratification approach as an aid to augment in the setting of a global public health crises, namely infectious diseases. Pharmaceutical companies have limited interest in budgeting drug discovery for niche markets and therapies with limited market value. Computational drug repositioning with modern clinical trial designs offer a comprehensive approach to address such challenges. In this work, we applied open access biomedical data, bioinformatics tools and methods to compile and interpret complex data to prioritize potential targets and recommend therapies for NiV, an emerging infectious disease. While we identified several indications, these drugs are not ready for immediate clinical use for NiV and need further experimental and clinical studies before use. In addition to pharmacological and biochemical annotations based filtering, the drugs can be filtered for approval status, availability, and cost as drug distribution varies across global markets. However, such filtering has not been considered in the study and drugs approved and available in the United States of America may or may not available with an approval in other countries or vice versa. In this study, we have only explored computational drug repositioning; other strategies including target-driven drug discovery is another strategy that may help to find a therapy for NiV infection. Several, well characterized host-pathogen interaction suggests targets including STAT1, Ephrins and ANP32B could be suitable for structure-based drug discovery and need further studies. Designing RNA interference molecules [105, 127] or peptide drugs to target these interactions may be useful to develop anti-NiV therapies [28, 32, 52, 78, 81, 82, 84, 89, 97, 128–168].

### From data to drugs with clinical evidence: Adaptive and n-of-1 clinical trials in the context of repositioned drugs targeting NiV

While several clinical trial design methods have been reported in the medical literature, randomized controlled trials (RCT) are considered to be the gold standard for gathering evidence for clinical use (See Kullo et.al and Jouni et.al; also see: https://clinicaltrials.gov/ct2/show/NCT01936675 for a clinical trial that assessed how genetic information might improve assessment of heart attack risk [169, 170]). However, clinical trial designs are evolving. Several new clinical trial design methods have been introduced over the last few decades (adaptive trial and n-of-1 trial)[171, 172]. These trials were primarily designed to augment RCT methods to develop evidence for personalized clinical modalities and interventions in the setting of precision medicine.

An adaptive trial is a clinical trial to evaluate the efficacy of a medical device or intervention by observing patient outcomes within a pre-defined schedule; the trial could be “adapted” to the observations from the intermediate endpoints. For example, given three therapies (A, B, and C) and a placebo (P), an adaptive trial can be designed with two intermediate endpoints (EP1 and EP2) and the end stage of a phase III trial. Therapies that meet EP1 (B and C) will proceed to EP2. If only one therapy is effective at EP2; only this therapy and placebo will proceed to the final phase of the trial. Adaptive trials are currently being evaluated as a trial design for various infectious diseases including HIV and Ebola [173–177].

The n-of-1 trial involves measuring a single patient repeatedly over time, while introducing different therapies (these could be multiple active therapies [178] or comparing therapy to placebo [179]). In addition to individual-level analyses to understand therapeutic effectiveness for a particular patient, several N-of-1 trials can be statistically aggregated to understand differences in effectiveness across patients, or provide population-level estimates [180]. Despite their benefits and alignment with personalized medicine approaches, N-of-1s present challenges in the context of NiV infection. NiV infection is associated with high mortality rate and complex outcomes including myocarditis, neurological complications and ARDS. These symptoms often appear as a singular presentation, and present in patients in different combinations; this makes it difficult to capture end points in the continuous fashion required by N-of-1 approaches. Delineating the clinical symptoms as endpoints would be key to developing a trial design strategy. A trial design that can accommodate features like survival analyses with time-to-event as an outcome, should also be explored. Furthermore, the strongest form of n-of-1 trials not only requires a baseline phase (without treatment) and treatment phase (where you introduce the drug) but also requires treatment withdrawal and washout periods to observe the effect before reintroducing the treatment. If a positive effect of the drug is observed, withdrawal may not be possible on ethical grounds, and washout periods without any treatments may also harm patient safety. Given these scenarios, other forms of single-subject designs that do not require treatment withdrawal (such as multiple baseline designs), or a baseline phase (such as alternating treatment designs) could be considered. To improve confidence in results from these trial designs, randomization could be incorporated [172, 181, 182]. To summarize, we do not endorse one method over the other but discussed various options of clinical trial design which follow personalized medicine approaches, and should be explored for infectious disease and multiple therapeutic options in the context of drug repositioning candidates.

## Limitations

Our research study has several limitations. Our data is based on infection induced in HUVEC cells. HUVEC cells may reflect differential gene expression from an affected tissue in a disease of interest. In the absence of patient derived genomic or transcriptomic data sets, the drug lists are less reliable for human testing without any additional experimental evidences. We have very low sample size and the gene signature may fluctuate to a higher or lower number of genes based on the sample size, statistical tests, and multiple testing correction methods. The sample size does not meet the biological replication criteria of a minimum of three samples. Our drug repositioning algorithms are predictive in nature and do not endorse immediate clinical use without further pre-clinical and clinical studies. Off-label prescription and recommendations would need additional comparative effectiveness trials for these drugs. Also, as a research study, the list of drugs compiled in the results section needs further evaluation in preclinical validation models and needs to be tested in patient cohorts using different clinical trial modalities including comparative effectiveness trials. Various factors drive the success of statistical validity of the predictive frameworks including availability and quality of data, choice of analytical algorithm, orthogonal evidence, and statistical methods that accounts for a large number of data points. We restrict the results, interpretation and discussions on drugs that are in DrugBank; however, other compounds may have superior anti-NiV activity. These compounds would need extensive, multi-year trials, and toxicity profiling before clinical testing and should not be used in clinical setting based on the predictive evidences.

## Future work

The gene signature used in this study is based on HUVEC cells, a cell line model for endothelial cells. This could be considered as a surrogate for infection as genes has been shown to express in a robust fashion across multiple tissues in the setting of human diseases. However, a cell line is identical as how infection express in the setting of the human body with multiple cell lines, tissues, organ systems. Also, the cell lines are a simplified representation of complex human disease physiology, where many patients could have several comorbidities and perturbed multiple factors including medications, environment, diet etc. These factors are not considered in the context of our study. Further, an ideal gene signature could be derived by comparing transcriptome extracted from whole blood of patients with NiV infection and age, gender-matched controls with no infection [183]. For example, we have performed similar analyses in diseases like peripheral arterial diseases and later used for drug repositioning investigations [106]. Further follow-up studies including biochemical testing and clinical trials such as adaptive and N-of-1 approaches are also needed.

## Conclusions

Drug repositioning could be a drug identification strategy in the absence of a ready to prescribe, FDA-approved therapies, in the context of an emerging virus with a high mortality rate in a short turnaround time. In this proof of concept study, we explored the application of computational drug repositioning and chemogenomic enrichment analyses using host-virus transcriptome data. Specifically, we performed computational drug matching searches to find drugs that can reverse the virus induced gene signature on an endothelial cell line in the setting of NiV infection. Briefly, translational bioinformatics approaches with epidemiology efforts may help to accelerate the discovery of affordable therapeutic options for public health crises including emergence of infectious diseases.

## Acknowledgements

KS would like to thank Dr. Anoop Kumar A S, MBBS, DM (Critical Care Medicine, Baby Memorial Hospital, Kozhikode (Calicut), Kerala, India; Dr. Paulo Varghese Akkara, MBBS, MD, DM, DNB, DTCD, Department of Pulmonary Medicine, Government Medical College, Kozhikode (Calicut), Kerala, India, Dr. Altaf Ali, MBBS, MD, Government Medical College, Manjeri, Kerala and members of Indian Medical Association Task Force For Nipah Virus, Kerala. Design, development and analytics of RepurposeDB are supported by the following National Institutes of Health (NIH) grants: National Institute of Diabetes and Digestive and Kidney Diseases (NIDDK, R01-DK098242-03); Illuminating the Druggable Genome (IDG), Knowledge Management Center sponsored by NIH Common Fund, National Cancer Institute (NCI, U54-CA189201-02); and Clinical and Translational Science Award (CTSA) by National Center for Advancing Translational Sciences (NCATS, UL1TR000067). Swiss Institute of Bioinformatics’ International Resource Innovation Award supports the update of RepurposeDB.

## References

1. Eaton BT, Broder CC, Middleton D, Wang LF: Hendra and Nipah viruses: different and dangerous. Nat Rev Microbiol 2006, 4(1):23–35.

2. Weatherman S, Feldmann H, de Wit E: Transmission of henipaviruses. Curr Opin Virol 2018, 28:7–11.

3. Williamson MM, Torres-Velez FJ: Henipavirus: a review of laboratory animal pathology. Vet Pathol 2010, 47(5):871–880.

4. Wild TF: Henipaviruses: a new family of emerging Paramyxoviruses. Pathol Biol (Paris) 2009, 57(2):188–196.

5. Fogarty R, Halpin K, Hyatt AD, Daszak P, Mungall BA: Henipavirus susceptibility to environmental variables. Virus Res 2008, 132(1-2):140–144.

6. Epstein JH, Prakash V, Smith CS, Daszak P, McLaughlin AB, Meehan G, Field HE, Cunningham AA: Henipavirus infection in fruit bats (Pteropus giganteus), India. Emerg InfectDis 2008, 14(8):1309–1311.

7. Field HE, Mackenzie JS, Daszak P: Henipaviruses: emerging paramyxoviruses associated with fruit bats. Curr Top Microbiol Immunol 2007, 315:133–159.

8. Mackenzie JS, Field HE: Emerging encephalitogenic viruses: lyssaviruses and henipaviruses transmitted by frugivorous bats. Arch Virol Suppl 2004(18):97–111.

9. Mackenzie JS, Field HE, Guyatt KJ: Managing emerging diseases borne by fruit bats (flying foxes), with particular reference to henipaviruses and Australian bat lyssavirus. JAppl Microbiol 2003, 94 Suppl:59S–69S.

10. Ang BSP, Lim TCC, Wang L: Nipah Virus Infection. J Clin Microbiol 2018.

11. Mahy BW, Brown CC: Emerging zoonoses: crossing the species barrier. Rev Sci Tech 2000, 19(1):33–40.

12. Centers for Disease C, Prevention: Update: outbreak of Nipah virus--Malaysia and Singapore, 1999. MMWR Morb Mortal Wkly Rep 1999, 48(16):335–337.

13. Chua KB, Goh KJ, Wong KT, Kamarulzaman A, Tan PS, Ksiazek TG, Zaki SR, Paul G, Lam SK, Tan CT: Fatal encephalitis due to Nipah virus among pig-farmers in Malaysia. Lancet 1999, 354(9186):1257–1259.

14. Farrar JJ: Nipah-virus encephalitis--investigation of a new infection. Lancet 1999, 354(9186):1222–1223.

15. Goh KJ, Tan CT, Chew NK, Tan PS, Kamarulzaman A, Sarji SA, Wong KT, Abdullah BJ, Chua KB, Lam SK: Clinical features of Nipah virus encephalitis among pig farmers in Malaysia. N Engl J Med 2000, 342(17):1229–1235.

16. Chong HT, Kunjapan SR, Thayaparan T, Tong J, Petharunam V, Jusoh MR, Tan CT: Nipah encephalitis outbreak in Malaysia, clinical features in patients from Seremban. Can J Neurol Sci 2002, 29(1):83–87.

17. Lim CC, Lee WL, Leo YS, Lee KE, Chan KP, Ling AE, Oh H, Auchus AP, Paton NI, Hui F et al: Late clinical and magnetic resonance imaging follow up of Nipah virus infection. J Neurol Neurosurg Psychiatry 2003, 74(1):131–133.

18. Williams JV: The clinical presentation and outcomes of children infected with newly identified respiratory tract viruses. Infect Dis Clin North Am 2005, 19(3):569–584.

19. Hossain MJ, Gurley ES, Montgomery JM, Bell M, Carroll DS, Hsu VP, Formenty P, Croisier A, Bertherat E, Faiz MA et al: Clinical presentation of nipah virus infection in Bangladesh. Clin Infect Dis 2008, 46(7):977–984.

20. Rockx B, Brining D, Kramer J, Callison J, Ebihara H, Mansfield K, Feldmann H: Clinical outcome of henipavirus infection in hamsters is determined by the route and dose of infection. J Virol 2011, 85(15):7658–7671.

21. Wong KT, Tan CT: Clinical and pathological manifestations of human henipavirus infection. Curr Top Microbiol Immunol 2012, 359:95–104.

22. Dups J, Middleton D, Long F, Arkinstall R, Marsh GA, Wang LF: Subclinical infection without encephalitis in mice following intranasal exposure to Nipah virus-Malaysia and Nipah virus-Bangladesh. Virol J 2014, 11:102.

23. Teng CL, Zuhanariah MN, Ng CS, Goh CC: Bibliography of clinical research in malaysia: methods and brief results. Med J Malaysia 2014, 69 Suppl A:4–7.

24. Kulkarni DD, Tosh C, Venkatesh G, Senthil Kumar D: Nipah virus infection: current scenario. Indian J Virol 2013, 24(3):398–408.

25. Wong KT, Shieh WJ, Kumar S, Norain K, Abdullah W, Guarner J, Goldsmith CS, Chua KB, Lam SK, Tan CT et al: Nipah virus infection: pathology and pathogenesis of an emerging paramyxoviral zoonosis. Am J Pathol 2002, 161(6):2153–2167.

26. Chan YP, Chua KB, Koh CL, Lim ME, Lam SK: Complete nucleotide sequences of Nipah virus isolates from Malaysia. J Gen Virol 2001, 82(Pt 9):2151–2155.

27. Griot C, Vandevelde M, Schobesberger M, Zurbriggen A: Canine distemper, a re-emerging morbillivirus with complex neuropathogenic mechanisms. Anim Health Res Rev 2003, 4(1):1–10.

28. Rodriguez JJ, Cruz CD, Horvath CM: Identification of the nuclear export signal and STAT-binding domains of the Nipah virus V protein reveals mechanisms underlying interferon evasion. J Virol 2004, 78(10):5358–5367.

29. Ciancanelli MJ, Volchkova VA, Shaw ML, Volchkov VE, Basler CF: Nipah virus sequesters inactive STAT1 in the nucleus via a P gene-encoded mechanism. J Virol 2009, 83(16):7828–7841.

30. Smith EC, Popa A, Chang A, Masante C, Dutch RE: Viral entry mechanisms: the increasing diversity of paramyxovirus entry. FEBS J 2009, 276(24):7217–7227.

31. Aguilar HC, Lee B: Emerging paramyxoviruses: molecular mechanisms and antiviral strategies. Expert Rev Mol Med 2011, 13:e6.

32. Chen W, Streaker ED, Russ DE, Feng Y, Prabakaran P, Dimitrov DS: Characterization of germline antibody libraries from human umbilical cord blood and selection of monoclonal antibodies to viral envelope glycoproteins: Implications for mechanisms of immune evasion and design of vaccine immunogens. Biochem Biophys Res Commun 2012, 417(4):1164–1169.

33. Liu Q, Stone JA, Bradel-Tretheway B, Dabundo J, Benavides Montano JA, Santos-Montanez J, Biering SB, Nicola AV, Iorio RM, Lu X et al: Unraveling a three-step spatiotemporal mechanism of triggering of receptor-induced Nipah virus fusion and cell entry. PLoSPathog 2013, 9(11):e1003770.

34. Chan YP, Koh CL, Lam SK, Wang LF: Mapping of domains responsible for nucleocapsid protein-phosphoprotein interaction of Henipaviruses. J Gen Virol 2004, 85(Pt 6):1675–1684.

35. Ludlow LE, Lo MK, Rodriguez JJ, Rota PA, Horvath CM: Henipavirus V protein association with Polo-like kinase reveals functional overlap with STAT1 binding and interferon evasion. J Virol 2008, 82(13):6259–6271.

36. Williams A, Henao-Mejia J, Harman CC, Flavell RA: miR-181 and metabolic regulation in the immune system. Cold Spring Harb Symp Quant Biol 2013, 78:223–230.

37. Henao-Mejia J, Williams A, Goff LA, Staron M, Licona-Limon P, Kaech SM, Nakayama M, Rinn JL, Flavell RA: The microRNA miR-181 is a critical cellular metabolic rheostat essential for NKT cell ontogenesis and lymphocyte development and homeostasis. Immunity 2013, 38(5):984–997.

38. Hutchison ER, Kawamoto EM, Taub DD, Lal A, Abdelmohsen K, Zhang Y, Wood WH, 3rd, Lehrmann E, Camandola S, Becker KG et al: Evidence for miR-181 involvement in neuroinflammatory responses of astrocytes. Glia 2013, 61(7):1018–1028.

39. Foo CH, Rootes CL, Cowley K, Marsh GA, Gould CM, Deffrasnes C, Cowled CJ, Klein R, Riddell SJ, Middleton D et al: Dual microRNA Screens Reveal That the Immune-Responsive miR-181 Promotes Henipavirus Entry and Cell-Cell Fusion. PLoS Pathog 2016, 12(10):e1005974.

40. Al-Obaidi MMJ, Bahadoran A, Wang SM, Manikam R, Raju CS, Sekaran SD: Disruption of the blood brain barrier is vital property of neurotropic viral infection of the central nervous system. Acta Virol 2018, 62(1):16–27.

41. Bauer A, Neumann S, Karger A, Henning AK, Maisner A, Lamp B, Dietzel E, Kwasnitschka L, Balkema-Buschmann A, Keil GM et al: ANP32B is a nuclear target of henipavirus M proteins. PLoS One 2014, 9(5):e97233.

42. Gupta M, Lo MK, Spiropoulou CF: Activation and cell death in human dendritic cells infected with Nipah virus. Virology 2013, 441(1):49–56.

43. Xu K, Broder CC, Nikolov DB: Ephrin-B2 and ephrin-B3 as functional henipavirus receptors. Semin Cell Dev Biol 2012, 23(1):116–123.

44. Lee B, Pernet O, Ahmed AA, Zeltina A, Beaty SM, Bowden TA: Molecular recognition of human ephrinB2 cell surface receptor by an emergent African henipavirus. Proc Natl Acad Sci U S A 2015, 112(17):E2156–2165.

45. Sugai A, Sato H, Yoneda M, Kai C: Gene end-like sequences within the 3’ noncoding region of the Nipah virus genome attenuate viral gene transcription. Virology 2017, 508:36–44.

46. Zhang G, Cowled C, Shi Z, Huang Z, Bishop-Lilly KA, Fang X, Wynne JW, Xiong Z, Baker ML, Zhao W et al: Comparative analysis of bat genomes provides insight into the evolution of flight and immunity. Science 2013, 339(6118):456–460.

47. Sleeman K, Bankamp B, Hummel KB, Lo MK, Bellini WJ, Rota PA: The C, V and W proteins of Nipah virus inhibit minigenome replication. J Gen Virol 2008, 89(Pt 5):1300–1308.

48. Habjan M, Andersson I, Klingstrom J, Schumann M, Martin A, Zimmermann P, Wagner V, Pichlmair A, Schneider U, Muhlberger E et al: Processing of genome 5’ termini as a strategy of negative-strand RNA viruses to avoid RIG-I-dependent interferon induction. PLoS One 2008, 3(4):e2032.

49. Freiberg A, Dolores LK, Enterlein S, Flick R: Establishment and characterization of plasmid-driven minigenome rescue systems for Nipah virus: RNA polymerase I- and T7-catalyzed generation of functional paramyxoviral RNA. Virology 2008, 370(1):33–44.

50. Halpin K, Bankamp B, Harcourt BH, Bellini WJ, Rota PA: Nipah virus conforms to the rule of six in a minigenome replication assay. J Gen Virol 2004, 85(Pt 3):701–707.

51. Chua KB, Wang LF, Lam SK, Eaton BT: Full length genome sequence of Tioman virus, a novel paramyxovirus in the genus Rubulavirus isolated from fruit bats in Malaysia. Arch Virol 2002, 147(7):1323–1348.

52. Wang LF, Yu M, Hansson E, Pritchard LI, Shiell B, Michalski WP, Eaton BT: The exceptionally large genome of Hendra virus: support for creation of a new genus within the family Paramyxoviridae. J Virol 2000, 74(21):9972–9979.

53. Snell NJ: Ribavirin therapy for Nipah virus infection. J Virol 2004, 78(18):10211.

54. Guillaume V, Contamin H, Loth P, Grosjean I, Courbot MC, Deubel V, Buckland R, Wild TF: Antibody prophylaxis and therapy against Nipah virus infection in hamsters. J Virol 2006, 80(4):1972–1978.

55. Porotto M, Orefice G, Yokoyama CC, Mungall BA, Realubit R, Sganga ML, Aljofan M, Whitt M, Glickman F, Moscona A: Simulating henipavirus multicycle replication in a screening assay leads to identification of a promising candidate for therapy. J Virol 2009, 83(10):5148–5155.

56. Pattabhi S, Wilkins CR, Dong R, Knoll ML, Posakony J, Kaiser S, Mire CE, Wang ML, Ireton RC, Geisbert TW et al: Targeting Innate Immunity for Antiviral Therapy through Small Molecule Agonists of the RLR Pathway. J Virol 2015, 90(5):2372–2387.

57. Mire CE, Satterfield BA, Geisbert JB, Agans KN, Borisevich V, Yan L, Chan YP, Cross RW, Fenton KA, Broder CC et al: Pathogenic Differences between Nipah Virus Bangladesh and Malaysia Strains in Primates: Implications for Antibody Therapy. Sci Rep 2016, 6:30916.

58. Louz D, Bergmans HE, Loos BP, Hoeben RC: Cross-species transfer of viruses: implications for the use of viral vectors in biomedical research, gene therapy and as live-virus vaccines. J Gene Med 2005, 7(10):1263–1274.

59. Mungall BA, Middleton D, Crameri G, Bingham J, Halpin K, Russell G, Green D, McEachern J, Pritchard LI, Eaton BT et al: Feline model of acute nipah virus infection and protection with a soluble glycoprotein-based subunit vaccine. J Virol 2006, 80(24):12293–12302.

60. Weingartl HM, Berhane Y, Caswell JL, Loosmore S, Audonnet JC, Roth JA, Czub M: Recombinant nipah virus vaccines protect pigs against challenge. J Virol 2006, 80(16):7929–7938.

61. Bowden TA, Crispin M, Harvey DJ, Aricescu AR, Grimes JM, Jones EY, Stuart DI: Crystal structure and carbohydrate analysis of Nipah virus attachment glycoprotein: a template for antiviral and vaccine design. J Virol 2008, 82(23):11628–11636.

62. McEachern JA, Bingham J, Crameri G, Green DJ, Hancock TJ, Middleton D, Feng YR, Broder CC, Wang LF, Bossart KN: A recombinant subunit vaccine formulation protects against lethal Nipah virus challenge in cats. Vaccine 2008, 26(31):3842–3852.

63. Chattopadhyay A, Rose JK: Complementing defective viruses that express separate paramyxovirus glycoproteins provide a new vaccine vector approach. J Virol 2011, 85(5):2004–2011.

64. Du L, Zhang X, Liu J, Jiang S: Protocol for recombinant RBD-based SARS vaccines: protein preparation, animal vaccination and neutralization detection. J Vis Exp 2011(51).

65. Pallister J, Middleton D, Wang LF, Klein R, Haining J, Robinson R, Yamada M, White J, Payne J, Feng YR et al: A recombinant Hendra virus G glycoprotein-based subunit vaccine protects ferrets from lethal Hendra virus challenge. Vaccine 2011, 29(34):5623–5630.

66. Walpita P, Barr J, Sherman M, Basler CF, Wang L: Vaccine potential of Nipah viruslike particles. PLoS One 2011, 6(4):e18437.

67. Bossart KN, Rockx B, Feldmann F, Brining D, Scott D, LaCasse R, Geisbert JB, Feng YR, Chan YP, Hickey AC et al: A Hendra virus G glycoprotein subunit vaccine protects African green monkeys from Nipah virus challenge. Sci Transl Med 2012, 4(146):146ra107.

68. Jiang S, Lu L, Liu Q, Xu W, Du L: Receptor-binding domains of spike proteins of emerging or re-emerging viruses as targets for development of antiviral vaccines. Emerg Microbes Infect 2012, 1(8):e13.

69. Kong D, Wen Z, Su H, Ge J, Chen W, Wang X, Wu C, Yang C, Chen H, Bu Z: Newcastle disease virus-vectored Nipah encephalitis vaccines induce B and T cell responses in mice and long-lasting neutralizing antibodies in pigs. Virology 2012, 432(2):327–335.

70. von Messling V, Cattaneo R: Virology. A henipavirus vaccine in sight. Science 2012, 337(6095):651–652.

71. Broder CC, Xu K, Nikolov DB, Zhu Z, Dimitrov DS, Middleton D, Pallister J, Geisbert TW, Bossart KN, Wang LF: A treatment for and vaccine against the deadly Hendra and Nipah viruses. Antiviral Res 2013, 100(1):8–13.

72. Mire CE, Versteeg KM, Cross RW, Agans KN, Fenton KA, Whitt MA, Geisbert TW: Single injection recombinant vesicular stomatitis virus vaccines protect ferrets against lethal Nipah virus disease. Virol J 2013, 10:353.

73. Pallister JA, Klein R, Arkinstall R, Haining J, Long F, White JR, Payne J, Feng YR, Wang LF, Broder CC et al: Vaccination of ferrets with a recombinant G glycoprotein subunit vaccine provides protection against Nipah virus disease for over 12 months. Virol J 2013, 10:237.

74. Ploquin A, Szecsi J, Mathieu C, Guillaume V, Barateau V, Ong KC, Wong KT, Cosset FL, Horvat B, Salvetti A: Protection against henipavirus infection by use of recombinant adeno-associated virus-vector vaccines. J Infect Dis 2013, 207(3):469–478.

75. Yoneda M, Georges-Courbot MC, Ikeda F, Ishii M, Nagata N, Jacquot F, Raoul H, Sato H, Kai C: Recombinant measles virus vaccine expressing the Nipah virus glycoprotein protects against lethal Nipah virus challenge. PLoS One 2013, 8(3):e58414.

76. DeBuysscher BL, Scott D, Marzi A, Prescott J, Feldmann H: Single-dose live-attenuated Nipah virus vaccines confer complete protection by eliciting antibodies directed against surface glycoproteins. Vaccine 2014, 32(22):2637–2644.

77. Lo MK, Bird BH, Chattopadhyay A, Drew CP, Martin BE, Coleman JD, Rose JK, Nichol ST, Spiropoulou CF: Single-dose replication-defective VSV-based Nipah virus vaccines provide protection from lethal challenge in Syrian hamsters. Antiviral Res 2014, 101:26–29.

78. Sakib MS, Islam MR, Hasan AK, Nabi AH: Prediction of epitope-based peptides for the utility of vaccine development from fusion and glycoprotein of nipah virus using in silico approach. Adv Bioinformatics 2014, 2014:402–492.

79. Sun J, Wei Y, Rauf A, Zhang Y, Ma Y, Zhang X, Shilo K, Yu Q, Saif YM, Lu X et al: Methyltransferase-defective avian metapneumovirus vaccines provide complete protection against challenge with the homologous Colorado strain and the heterologous Minnesota strain. J Virol 2014, 88(21):12348–12363.

80. Yoneda M: [Study of pathogenicity of Nipah virus and its vaccine development]. Uirusu 2014, 64(1):105–112.

81. Ali MT, Morshed MM, Hassan F: A Computational Approach for Designing a Universal Epitope-Based Peptide Vaccine Against Nipah Virus. Interdiscip Sci 2015, 7(2):177–185.

82. Kamthania M, Sharma DK: Screening and structure-based modeling of T-cell epitopes of Nipah virus proteome: an immunoinformatic approach for designing peptide-based vaccine. 3 Biotech 2015, 5(6):877–882.

83. Kurup D, Wirblich C, Feldmann H, Marzi A, Schnell MJ: Rhabdovirus-based vaccine platforms against henipaviruses. J Virol 2015, 89(1):144–154.

84. Oyarzun P, Ellis JJ, Gonzalez-Galarza FF, Jones AR, Middleton D, Boden M, Kobe B: A bioinformatics tool for epitope-based vaccine design that accounts for human ethnic diversity: application to emerging infectious diseases. Vaccine 2015, 33(10):1267–1273.

85. Prescott J, DeBuysscher BL, Feldmann F, Gardner DJ, Haddock E, Martellaro C, Scott D, Feldmann H: Single-dose live-attenuated vesicular stomatitis virus-based vaccine protects African green monkeys from Nipah virus disease. Vaccine 2015, 33(24):2823–2829.

86. Broder CC, Weir DL, Reid PA: Hendra virus and Nipah virus animal vaccines. Vaccine 2016, 34(30):3525–3534.

87. DeBuysscher BL, Scott D, Thomas T, Feldmann H, Prescott J: Peri-exposure protection against Nipah virus disease using a single-dose recombinant vesicular stomatitis virus-based vaccine. NPJ Vaccines 2016, 1.

88. Guillaume-Vasselin V, Lemaitre L, Dhondt KP, Tedeschi L, Poulard A, Charreyre C, Horvat B: Protection from Hendra virus infection with Canarypox recombinant vaccine. NPJ Vaccines 2016, 1:16003.

89. Parvege MM, Rahman M, Nibir YM, Hossain MS: Two highly similar LAEDDTNAQKT and LTDKIGTEI epitopes in G glycoprotein may be useful for effective epitope based vaccine design against pathogenic Henipavirus. Comput Biol Chem 2016, 61:270–280.

90. Satterfield BA, Dawes BE, Milligan GN: Status of vaccine research and development of vaccines for Nipah virus. Vaccine 2016, 34(26):2971–2975.

91. Whitt MA, Geisbert TW, Mire CE: Single-Vector, Single-Injection Recombinant Vesicular Stomatitis Virus Vaccines Against High-Containment Viruses. Methods Mol Biol 2016, 1403:295–311.

92. Zhang Y, Sun J, Wei Y, Li J: A Reverse Genetics Approach for the Design of Methyltransferase-Defective Live Attenuated Avian Metapneumovirus Vaccines. Methods Mol Biol 2016, 1404:103–121.

93. Tackling Nipah virus: pound2.36 million grant awarded to Pirbright to create vaccine. Vet Rec 2017, 181(10):253.

94. Butler D: Billion-dollar project aims to prep vaccines before epidemics hit. Nature 2017, 541(7638):444–445.

95. Ewer K, Sebastian S, Spencer AJ, Gilbert S, Hill AVS, Lambe T: Chimpanzee adenoviral vectors as vaccines for outbreak pathogens. Hum Vaccin Immunother 2017, 13(12):3020–3032.

96. Plotkin SA: Vaccines for epidemic infections and the role of CEPI. Hum Vaccin Immunother 2017, 13(12):2755–2762.

97. Saha CK, Mahbub Hasan M, Saddam Hossain M, Asraful Jahan M, Azad AK: In silico identification and characterization of common epitope-based peptide vaccine for Nipah and Hendra viruses. Asian PacJ Trop Med 2017, 10(6):529–538.

98. Walpita P, Cong Y, Jahrling PB, Rojas O, Postnikova E, Yu S, Johns L, Holbrook MR: A VLP-based vaccine provides complete protection against Nipah virus challenge following multiple-dose or single-dose vaccination schedules in a hamster model. NPJ Vaccines 2017, 2:21.

99. Chong HT, Kamarulzaman A, Tan CT, Goh KJ, Thayaparan T, Kunjapan SR, Chew NK, Chua KB, Lam SK: Treatment of acute Nipah encephalitis with ribavirin. Ann Neurol 2001, 49(6):810–813.

100. Snell NJ: Ribavirin--current status of a broad spectrum antiviral agent. Expert Opin Pharmacother 2001, 2(8):1317–1324.

101. Georges-Courbot MC, Contamin H, Faure C, Loth P, Baize S, Leyssen P, Neyts J, Deubel V: Poly(I)-poly(C12U) but not ribavirin prevents death in a hamster model of Nipah virus infection. Antimicrob Agents Chemother 2006, 50(5):1768–1772.

102. Freiberg AN, Worthy MN, Lee B, Holbrook MR: Combined chloroquine and ribavirin treatment does not prevent death in a hamster model of Nipah and Hendra virus infection. J Gen Virol 2010, 91(Pt 3):765–772.

103. Rockx B, Bossart KN, Feldmann F, Geisbert JB, Hickey AC, Brining D, Callison J, Safronetz D, Marzi A, Kercher L et al: A novel model of lethal Hendra virus infection in African green monkeys and the effectiveness of ribavirin treatment. J Virol 2010, 84(19):9831–9839.

104. Dawes BE, Kalveram B, Ikegami T, Juelich T, Smith JK, Zhang L, Park A, Lee B, Komeno T, Furuta Y et al: Favipiravir (T-705) protects against Nipah virus infection in the hamster model. Sci Rep 2018, 8(1):7604.

105. Mungall BA, Schopman NC, Lambeth LS, Doran TJ: Inhibition of Henipavirus infection by RNA interference. Antiviral Res 2008, 80(3):324–331.

106. Chu LH, Annex BH, Popel AS: Computational drug repositioning for peripheral arterial disease: prediction of anti-inflammatory and pro-angiogenic therapeutics. Front Pharmacol 2015, 6:179.

107. Shameer K, Readhead B, Dudley JT: Computational and experimental advances in drug repositioning for accelerated therapeutic stratification. Curr Top Med Chem 2015, 15(1):5–20.

108. Shameer K, Glicksberg BS, Hodos R, Johnson KW, Badgeley MA, Readhead B, Tomlinson MS, O’Connor T, Miotto R, Kidd BA et al: Systematic analyses of drugs and disease indications in RepurposeDB reveal pharmacological, biological and epidemiological factors influencing drug repositioning. Brief Bioinform 2017.

109. Hodos RA, Kidd BA, Shameer K, Readhead BP, Dudley JT: In silico methods for drug repurposing and pharmacology. Wiley Interdiscip Rev Syst Biol Med 2016, 8(3):186–210.

110. Molineris I, Ala U, Provero P, Di Cunto F: Drug repositioning for orphan genetic diseases through Conserved Anticoexpressed Gene Clusters (CAGCs). BMC Bioinformatics 2013, 14:288.

111. Jahchan NS, Dudley JT, Mazur PK, Flores N, Yang D, Palmerton A, Zmoos AF, Vaka D, Tran KQ, Zhou M et al: A drug repositioning approach identifies tricyclic antidepressants as inhibitors of small cell lung cancer and other neuroendocrine tumors. Cancer Discov 2013, 3(12):1364–1377.

112. Sirota M, Dudley JT, Kim J, Chiang AP, Morgan AA, Sweet-Cordero A, Sage J, Butte AJ: Discovery and preclinical validation of drug indications using compendia of public gene expression data. Sci Transl Med 2011, 3(96):96ra77.

113. Sardana D, Zhu C, Zhang M, Gudivada RC, Yang L, Jegga AG: Drug repositioning for orphan diseases. Brief Bioinform 2011, 12(4):346–356.

114. Dudley JT, Deshpande T, Butte AJ: Exploiting drug-disease relationships for computational drug repositioning. Brief Bioinform 2011, 12(4):303–311.

115. Butte AJ, Chen R: Finding disease-related genomic experiments within an international repository: first steps in translational bioinformatics. AMIA Annu Symp Proc 2006:106–110.

116. Butte AJ: Translational bioinformatics: coming of age. J Am Med Inform Assoc 2008, 15(6):709–714.

117. Clough E, Barrett T: The Gene Expression Omnibus Database. Methods Mol Biol 2016, 1418:93–110.

118. Mathieu C, Guillaume V, Sabine A, Ong KC, Wong KT, Legras-Lachuer C, Horvat B: Lethal Nipah virus infection induces rapid overexpression of CXCL10. PLoS One 2012, 7(2):e32157.

119. Wong KT: Emerging and re-emerging epidemic encephalitis: a tale of two viruses. NeuropatholApplNeurobiol 2000, 26(4):313–318.

120. Magri A, Reilly R, Scalacci N, Radi M, Hunter M, Ripoll M, Patel AH, Castagnolo D: Rethinking the old antiviral drug moroxydine: Discovery of novel analogues as anti-hepatitis C virus (HCV) agents. Bioorg Med Chem Lett 2015, 25(22):5372–5376.

121. Fagerberg L, Hallstrom BM, Oksvold P, Kampf C, Djureinovic D, Odeberg J, Habuka M, Tahmasebpoor S, Danielsson A, Edlund K et al: Analysis of the human tissue-specific expression by genome-wide integration of transcriptomics and antibody-based proteomics. Mol Cell Proteomics 2014, 13(2):397–406.

122. van Hooij A, Boeters DM, Tjon Kon Fat EM, van den Eeden SJF, Corstjens P, van der Helm-van Mil AHM, Geluk A: Longitudinal IP-10 Serum Levels Are Associated with the Course of Disease Activity and Remission in Patients with Rheumatoid Arthritis. Clin Vaccine Immunol 2017, 24(8).

123. Sandborn WJ, Rutgeerts P, Colombel JF, Ghosh S, Petryka R, Sands BE, Mitra P, Luo A: Eldelumab [anti-interferon-gamma-inducible protein-10 antibody] Induction Therapy for Active Crohn’s Disease: a Randomised, Double-blind, Placebo-controlled Phase IIa Study. J Crohns Colitis 2017, 11(7):811–819.

124. Peters LA, Perrigoue J, Mortha A, Iuga A, Song WM, Neiman EM, Llewellyn SR, Di Narzo A, Kidd BA, Telesco SE et al: A functional genomics predictive network model identifies regulators of inflammatory bowel disease. Nat Genet 2017, 49(10):1437–1449.

125. Shameer K, Badgeley MA, Miotto R, Glicksberg BS, Morgan JW, Dudley JT: Translational bioinformatics in the era of real-time biomedical, health care and wellness data streams. Brief Bioinform 2017, 18(1):105–124.

126. Shameer K, Johnson KW, Glicksberg BS, Dudley JT, Sengupta PP: Machine learning in cardiovascular medicine: are we there yet? Heart 2018.

127. Stewart CR, Deffrasnes C, Foo CH, Bean AGD, Wang LF: A Functional Genomics Approach to Henipavirus Research: The Role of Nuclear Proteins, MicroRNAs and Immune Regulators in Infection and Disease. Curr Top Microbiol Immunol 2017.

128. Mathieu C, Porotto M, Figueira T, Horvat B, Moscona A: Fusion Inhibitory Lipopeptides Engineered for Prophylaxis of Nipah Virus in Primates. J Infect Dis 2018.

129. Mathieu C, Augusto MT, Niewiesk S, Horvat B, Palermo LM, Sanna G, Madeddu S, Huey D, Castanho MA, Porotto M et al: Broad spectrum antiviral activity for paramyxoviruses is modulated by biophysical properties of fusion inhibitory peptides. Sci Rep 2017, 7:43610.

130. Lam CW, AbuBakar S, Chang LY: Identification of the cell binding domain in Nipah virus G glycoprotein using a phage display system. J Virol Methods 2017, 243:1–9.

131. Augusto MT, Hollmann A, Porotto M, Moscona A, Santos NC: Antiviral Lipopeptide-Cell Membrane Interaction Is Influenced by PEG Linker Length. Molecules 2017, 22(7).

132. Yabukarski F, Leyrat C, Martinez N, Communie G, Ivanov I, Ribeiro EA, Jr., Buisson M, Gerard FC, Bourhis JM, Jensen MR et al: Ensemble Structure of the Highly Flexible Complex Formed between Vesicular Stomatitis Virus Unassembled Nucleoprotein and its Phosphoprotein Chaperone. J Mol Biol 2016, 428(13):2671–2694.

133. Wong JJ, Paterson RG, Lamb RA, Jardetzky TS: Structure and stabilization of the Hendra virus F glycoprotein in its prefusion form. Proc Natl Acad Sci U S A 2016, 113(4):1056–1061.

134. Uchida S, Sato H, Yoneda M, Kai C: Eukaryotic elongation factor 1-beta interacts with the 5’ untranslated region of the M gene of Nipah virus to promote mRNA translation. Arch Virol 2016, 161(9):2361–2368.

135. Stone JA, Vemulapati BM, Bradel-Tretheway B, Aguilar HC: Multiple Strategies Reveal a Bidentate Interaction between the Nipah Virus Attachment and Fusion Glycoproteins. J Virol 2016, 90(23):10762–10773.

136. Fischer K, dos Reis VP, Finke S, Sauerhering L, Stroh E, Karger A, Maisner A, Groschup MH, Diederich S, Balkema-Buschmann A: Expression, characterisation and antigenicity of a truncated Hendra virus attachment protein expressed in the protozoan host Leishmania tarentolae. J Virol Methods 2016, 228:48–54.

137. Weis M, Behner L, Binger T, Drexler JF, Drosten C, Maisner A: Fusion activity of African henipavirus F proteins with a naturally occurring start codon directly upstream of the signal peptide. Virus Res 2015, 201:85–93.

138. Mathieu C, Horvat B: Henipavirus pathogenesis and antiviral approaches. Expert Rev Anti Infect Ther 2015, 13(3):343–354.

139. Yabukarski F, Lawrence P, Tarbouriech N, Bourhis JM, Delaforge E, Jensen MR, Ruigrok RW, Blackledge M, Volchkov V, Jamin M: Structure of Nipah virus unassembled nucleoprotein in complex with its viral chaperone. Nat Struct Mol Biol 2014, 21(9):754–759.

140. Sun W, McCrory TS, Khaw WY, Petzing S, Myers T, Schmitt AP: Matrix proteins of Nipah and Hendra viruses interact with beta subunits of AP-3 complexes. J Virol 2014, 88(22):13099–13110.

141. Elshabrawy HA, Fan J, Haddad CS, Ratia K, Broder CC, Caffrey M, Prabhakar BS: Identification of a broad-spectrum antiviral small molecule against severe acute respiratory syndrome coronavirus and Ebola, Hendra, and Nipah viruses by using a novel high-throughput screening assay. J Virol 2014, 88(8):4353–4365.

142. Pessi A, Langella A, Capito E, Ghezzi S, Vicenzi E, Poli G, Ketas T, Mathieu C, Cortese R, Horvat B et al: A general strategy to endow natural fusion-protein-derived peptides with potent antiviral activity. PLoS One 2012, 7(5):e36833.

143. Dochow M, Krumm SA, Crowe JE, Jr., Moore ML, Plemper RK: Independent structural domains in paramyxovirus polymerase protein. J Biol Chem 2012, 287(9):6878–6891.

144. Chan YP, Lu M, Dutta S, Yan L, Barr J, Flora M, Feng YR, Xu K, Nikolov DB, Wang LF et al: Biochemical, conformational, and immunogenic analysis of soluble trimeric forms of henipavirus fusion glycoproteins. J Virol 2012, 86(21):11457–11471.

145. Talekar A, Pessi A, Porotto M: Infection of primary neurons mediated by nipah virus envelope proteins: role of host target cells in antiviral action. J Virol 2011, 85(16):8422–8426.

146. Porotto M, Devito I, Palmer SG, Jurgens EM, Yee JL, Yokoyama CC, Pessi A, Moscona A: Spring-loaded model revisited: paramyxovirus fusion requires engagement of a receptor binding protein beyond initial triggering of the fusion protein. J Virol 2011, 85(24):12867–12880.

147. Huang M, Sato H, Hagiwara K, Watanabe A, Sugai A, Ikeda F, Kozuka-Hata H, Oyama M, Yoneda M, Kai C: Determination of a phosphorylation site in Nipah virus nucleoprotein and its involvement in virus transcription. J Gen Virol 2011, 92(Pt 9):2133–2141.

148. Shameer K, Madan LL, Veeranna S, Gopal B, Sowdhamini R: PeptideMine--a webserver for the design of peptides for protein-peptide binding studies derived from protein-protein interactomes. BMC Bioinformatics 2010, 11:473.

149. Porotto M, Yokoyama CC, Palermo LM, Mungall B, Aljofan M, Cortese R, Pessi A, Moscona A: Viral entry inhibitors targeted to the membrane site of action. J Virol 2010, 84(13):6760–6768.

150. Porotto M, Rockx B, Yokoyama CC, Talekar A, Devito I, Palermo LM, Liu J, Cortese R, Lu M, Feldmann H et al: Inhibition of Nipah virus infection in vivo: targeting an early stage of paramyxovirus fusion activation during viral entry. PLoS Pathog 2010, 6(10):e1001168.

151. Pei Z, Bai Y, Schmitt AP: PIV5 M protein interaction with host protein angiomotin-like 1. Virology 2010, 397(1):155–166.

152. McGinnes LW, Pantua H, Laliberte JP, Gravel KA, Jain S, Morrison TG: Assembly and biological and immunological properties of Newcastle disease virus-like particles. J Virol 2010, 84(9):4513–4523.

153. Khetawat D, Broder CC: A functional henipavirus envelope glycoprotein pseudotyped lentivirus assay system. Virol J 2010, 7:312.

154. Aguilar HC, Aspericueta V, Robinson LR, Aanensen KE, Lee B: A quantitative and kinetic fusion protein-triggering assay can discern distinct steps in the nipah virus membrane fusion cascade. J Virol 2010, 84(16):8033–8041.

155. Lo MK, Harcourt BH, Mungall BA, Tamin A, Peeples ME, Bellini WJ, Rota PA: Determination of the henipavirus phosphoprotein gene mRNA editing frequencies and detection of the C, V and W proteins of Nipah virus in virus-infected cells. J Gen Virol 2009, 90(Pt 2):398–404.

156. Diederich S, Dietzel E, Maisner A: Nipah virus fusion protein: influence of cleavage site mutations on the cleavability by cathepsin L, trypsin and furin. Virus Res 2009, 145(2):300–306.

157. Aguilar HC, Ataman ZA, Aspericueta V, Fang AQ, Stroud M, Negrete OA, Kammerer RA, Lee B: A novel receptor-induced activation site in the Nipah virus attachment glycoprotein (G) involved in triggering the fusion glycoprotein (F). J Biol Chem 2009, 284(3):1628–1635.

158. Xiao C, Liu Y, Jiang Y, Magoffin DE, Guo H, Xuan H, Wang G, Wang LF, Tu C: Monoclonal antibodies against the nucleocapsid proteins of henipaviruses: production, epitope mapping and application in immunohistochemistry. Arch Virol 2008, 153(2):273–281.

159. Aljofan M, Porotto M, Moscona A, Mungall BA: Development and validation of a chemiluminescent immunodetection assay amenable to high throughput screening of antiviral drugs for Nipah and Hendra virus. J Virol Methods 2008, 149(1):12–19.

160. Porotto M, Carta P, Deng Y, Kellogg GE, Whitt M, Lu M, Mungall BA, Moscona A: Molecular determinants of antiviral potency of paramyxovirus entry inhibitors. J Virol 2007, 81(19):10567–10574.

161. Zhu Z, Dimitrov AS, Bossart KN, Crameri G, Bishop KA, Choudhry V, Mungall BA, Feng YR, Choudhary A, Zhang MY et al: Potent neutralization of Hendra and Nipah viruses by human monoclonal antibodies. J Virol 2006, 80(2):891–899.

162. Eshaghi M, Tan WS, Yusoff K: Identification of epitopes in the nucleocapsid protein of Nipah virus using a linear phage-displayed random peptide library. J Med Virol 2005, 75(1):147–152.

163. Bossart KN, Mungall BA, Crameri G, Wang LF, Eaton BT, Broder CC: Inhibition of Henipavirus fusion and infection by heptad-derived peptides of the Nipah virus fusion glycoprotein. Virol J 2005, 2:57.

164. Bossart KN, Crameri G, Dimitrov AS, Mungall BA, Feng YR, Patch JR, Choudhary A, Wang LF, Eaton BT, Broder CC: Receptor binding, fusion inhibition, and induction of cross-reactive neutralizing antibodies by a soluble G glycoprotein of Hendra virus. J Virol 2005, 79(11):6690–6702.

165. Xu Y, Gao S, Cole DK, Zhu J, Su N, Wang H, Gao GF, Rao Z: Basis for fusion inhibition by peptides: analysis of the heptad repeat regions of the fusion proteins from Nipah and Hendra viruses, newly emergent zoonotic paramyxoviruses. Biochem Biophys Res Commun 2004, 315(3):664–670.

166. Johansson K, Bourhis JM, Campanacci V, Cambillau C, Canard B, Longhi S: Crystal structure of the measles virus phosphoprotein domain responsible for the induced folding of the C-terminal domain of the nucleoprotein. J Biol Chem 2003, 278(45):44567–44573.

167. Shiell BJ, Beddome G, Michalski WP: Mass spectrometric identification and characterisation of the nucleocapsid protein of Menangle virus. J Virol Methods 2002, 102(1-2):27–35.

168. Bossart KN, Wang LF, Flora MN, Chua KB, Lam SK, Eaton BT, Broder CC: Membrane fusion tropism and heterotypic functional activities of the Nipah virus and Hendra virus envelope glycoproteins. J Virol 2002, 76(22):11186–11198.

169. Jouni H, Haddad RA, Marroush TS, Brown SA, Kruisselbrink TM, Austin EE, Shameer K, Behnken EM, Chaudhry R, Montori VM et al: Shared decision-making following disclosure of coronary heart disease genetic risk: results from a randomized clinical trial. JInvestig Med 2017, 65(3):681–688.

170. Kullo IJ, Jouni H, Austin EE, Brown SA, Kruisselbrink TM, Isseh IN, Haddad RA, Marroush TS, Shameer K, Olson JE et al: Incorporating a Genetic Risk Score Into Coronary Heart Disease Risk Estimates: Effect on Low-Density Lipoprotein Cholesterol Levels (the MI-GENES Clinical Trial). Circulation 2016, 133(12):1181–1188.

171. Kairalla JA, Coffey CS, Thomann MA, Muller KE: Adaptive trial designs: a review of barriers and opportunities. Trials 2012, 13:145.

172. Schork NJ: Personalized medicine: Time for one-person trials. Nature 2015, 520(7549):609–611.

173. Huskins WC, Fowler VG, Jr., Evans S: Adaptive Designs for Clinical Trials: Application to Healthcare Epidemiology Research. Clin Infect Dis 2018, 66(7):1140–1146.

174. Lanini S, Zumla A, Ioannidis JP, Di Caro A, Krishna S, Gostin L, Girardi E, Pletschette M, Strada G, Baritussio A et al: Are adaptive randomised trials or nonrandomised studies the best way to address the Ebola outbreak in west Africa? Lancet Infect Dis 2015, 15(6):738–745.

175. Cook J, Verheij T, van der Velden A, Goossens H, de Jong M, Beutels P, Little P, Butler C: Using an adaptive trial design for an infectious disease in primacy care - challenges in the design and set-up stages. Trials 2015, 16(Suppl 2):P211–P211.

176. Corey L, Nabel GJ, Dieffenbach C, Gilbert P, Haynes BF, Johnston M, Kublin J, Lane HC, Pantaleo G, Picker LJ et al: HIV-1 vaccines and adaptive trial designs. Sci Transl Med 2011, 3(79):79ps13.

177. Dodd LE, Proschan MA, Neuhaus J, Koopmeiners JS, Neaton J, Beigel JD, Barrett K, Lane HC, Davey RT, Jr.: Design of a Randomized Controlled Trial for Ebola Virus Disease Medical Countermeasures: PREVAIL II, the Ebola MCM Study. J Infect Dis 2016, 213(12):1906–1913.

178. Alemayehu C, Mitchell G, Aseffa A, Clavarino A, McGree J, Nikles J: A series of N-of-1 trials to assess the therapeutic interchangeability of two enalapril formulations in the treatment of hypertension in Addis Ababa, Ethiopia: study protocol for a randomized controlled trial. Trials 2017, 18(1):470.

179. Hackett A, Gillard J, Wilcken B: n of 1 trial for an ornithine transcarbamylase deficiency carrier. Mol Genet Metab 2008, 94(2):157–161.

180. Zucker DR, Ruthazer R, Schmid CH: Individual (N-of-1) trials can be combined to give population comparative treatment effect estimates: methodologic considerations. J Clin Epidemiol 2010, 63(12):1312–1323.

181. Zucker DR, Schmid CH, McIntosh MW, D’Agostino RB, Selker HP, Lau J: Combining single patient (N-of-1) trials to estimate population treatment effects and to evaluate individual patient responses to treatment. J Clin Epidemiol 1997, 50(4):401–410.

182. Lillie EO, Patay B, Diamant J, Issell B, Topol EJ, Schork NJ: The n-of-1 clinical trial: the ultimate strategy for individualizing medicine? Per Med 2011, 8(2):161–173.

183. Stachowiak B, Weingartl HM: Nipah virus infects specific subsets of porcine peripheral blood mononuclear cells. PLoS One 2012, 7(1):e30855.

184. Tobinick EL: The value of drug repositioning in the current pharmaceutical market. Drug News Perspect 2009, 22(2):119–125.

